# Efficacy of a bacteriophage cocktail targeting pathogenic bacteria associated with gastrointestinal dysfunction

**DOI:** 10.1101/2025.10.03.680221

**Authors:** Yuansong Li, Ashley E Franks, Steve Petrovski, Elisa L Hill-Yardin

## Abstract

The use of bacteriophage viruses that infect and kill bacteria is increasingly explored as an alternative to antibiotics for treatment of infections caused by opportunistic pathogens. Examples of opportunistic pathogens that disrupt the intestinal barrier and contribute to microbial imbalance in the gastrointestinal tract include *Klebsiella pneumoniae*, *Pseudomonas aeruginosa* and *Serratia marcescens*. This study evaluated the *in vitro* effectiveness of a bacteriophage cocktail composed of three host-specific phages: Lilla1 targeting *Klebsiella pneumoniae*, PAE1 targeting *Pseudomonas aeruginosa* and Smarc1 targeting *Serratia marcescens*. Bacterial growth was monitored by measuring optical density at 600 nm, and phage replication was quantified by plaque-forming unit assays. Each phage exhibited strong host specificity and produced significant reduction of growth of the corresponding bacterium while increasing in concentration during incubation. When applied as a cocktail, these three phages produced stronger and more sustained suppression of bacterial growth than single-phage treatments, both in individual cultures and in mixed cultures containing all three pathogens. In mixed cultures, continuous inhibition of bacterial growth persisted throughout a 24-hour incubation, and concentrations of Lilla1, PAE1 and Smarc1 increased significantly, indicating active replication and enhanced efficacy in combination. In single bacterial growth assays, bacterial populations that were initially suppressed showed regrowth after extended incubation. Regrowth manifested as a renewed increase in optical density several hours after initial decline and is consistent with selection of phage-resistant subpopulations, potentially arising from modification of surface receptors, phase variation or activation of adaptive defence systems. The observed biphasic response highlights the need for measures to limit microbial resistance to phage treatment, including expanded phage cocktails, adjusted dosing strategies and combined phage-antibiotic regimens. The results demonstrate that a three-phage cocktail can simultaneously inhibit multiple bacterial pathogens associated with gastrointestinal dysfunction and provide a basis for further *in vivo* evaluation and therapeutic development.

## Introduction

Previous studies report that gram-negative bacteria, including *Klebsiella pneumoniae* (*K. pneumoniae*), *Pseudomonas aeruginosa* (*P. aeruginosa*), and *Serratia marcescens* (*S. marcescens*), can contribute to increased intestinal epithelial permeability [1, 2]. Lipopolysaccharides (LPS) derived from gram-negative bacteria like *Escherichia coli*, *K. pneumoniae*, *P. aeruginosa*, and *S. marcescens* modulate intestinal epithelial permeability. These pathogens have also been demonstrated to increase gastrointestinal (GI) permeability *in vitro* as reported using human intestinal epithelial cell monolayers (Caco-2 cells) [2]. Such imbalances can trigger abnormal immune responses, irregular neuronal signalling, and GI symptoms [3, 4]. Increased GI mucosal permeability can allow microbiota to escape the intestinal lumen to invade host tissues and the blood stream and disrupt the microbial community profile within the GI tract [3, 5].Addressing dysregulated gut microbial populations may therefore be a promising approach for restoring impaired GI function.

*K. pneumoniae, P. aeruginosa*, and *S. marcescens* are gram-negative opportunistic pathogens. All three bacteria are implicated in respiratory diseases, including bronchopneumonia and lung abscesses (*K. pneumoniae*), chronic lung infections and severe pneumonia (*P. aeruginosa*), and pneumonia (*S. marcescens*) [6] [7] [8]. These pathogens are also implicated in GI dysfunction; for instance, *K. pneumoniae* infections in the GI tract can result in gastroenteritis, colitis, and liver abscesses, disrupting gut microbiota homeostasis and contributing to inflammatory bowel disease (IBD) [9]. *P. aeruginosa* can cause diarrhea, colitis, and severe conditions such as necrotizing enterocolitis in infants [10]. Similarly, *S. marcescens* has been associated with primary GI infections, such as gastroenteritis, particularly in neonates and immunocompromised individuals [11].

Current therapeutic approaches targeting pathogenic bacteria in the GI tract, particularly for restoring GI permeability, remain underdeveloped. While antibiotics have drastically improved public health and extended human lifespan, and treatments like probiotics and faecal microbiota transplantation (FMT) have demonstrated some success, the growing challenge of antibiotic resistance underscores the urgent need for alternative, non-antibiotic strategies that offer broader and more sustainable public health benefits [12–15]. However, these treatments have limitations including the potential for antibiotics to lead to drug resistance [16]. In addition, certain probiotics may pose risks to immunocompromised patients [17], and FMT carries the potential to transfer pathogens from donor to recipient [18, 19].

In addition to high specificity, phage offer a potential advantage in combating antimicrobial resistance [20]. Phage exhibit high specificity toward their bacterial hosts, which allows for precise targeting without disrupting beneficial microbiota. Phage therapy has been demonstrated to modify the GI environment by targeting intestinal pathogens [21]. Phage may also support the GI barrier by interacting with the mucosal surface, especially through binding to mucin, a key part of mucus that helps trap microbes. This happens when phage capsid proteins attach to the glycans on mucin, allowing phage to stay longer in the mucus and possibly act as an added defence at the mucosal layer [22, 23].

Here, we evaluated the *in vitro* efficacy of three phage strains; *K. pneumoniae* phage Lilla1, *P. aeruginosa* phage PAE1, and *S. marcescens* phage Smarc1 to inhibit the growth of their respective bacterial hosts.

## Materials and Methods

### Bacterial strains and culture

Three bacterial strains were used in this study: *Klebsiella pneumoniae*, *Pseudomonas aeruginosa* PAO9503, and *Serratia marcescens* Y21. All strains were obtained from the La Trobe University culture collection and stored at −80°C. Prior to experiments, the bacterial strains were cultured overnight on nutrient agar (NA) plates at 37°C (Table 1).

**Table 1.** Bacterial strains and target phage used.

| Bacterial Strains | Phage | Source |
| --- | --- | --- |
| *Klebsiella pneumoniae* | Lilla1 | La Trobe University Culture collection<br>(S. Petrovski) |
| *Pseudomonas aeruginosa* PAO9503 | PAE1 | La Trobe University Culture collection<br>(S. Petrovski) |
| *Serratia marcescens* Y21 | Smarc1 | La Trobe University Culture collection<br>(S. Petrovski) |

### Bacteriophage strains

Three bacteriophages, each specific to a single bacterial target strain, were used (i.e., Lilla1 for *K. pneumoniae*, PAE1 for *P. aeruginosa* PAO9503, and Smarc1 for *S. marcescens* Y21).

### Bacteriophage inoculation and filtration

Bacterial and phage strains were grown in 10ml nutrient broth (NB) and incubated at 37°C (as the optimal temperature to encourage growth of the pathogenic bacteria) overnight on a shaker. After phage inoculation, bacteria with cognate phage mixed cultures were centrifuged at 44,000rpm for 5min. The supernatant was filtered through a 0.22µm syringe filter and stored at 4°C as phage stock.

### Bacteriophage propagation and collection

For phage propagation, 300µl of phage stock was spread onto a bacterial lawned NA plate and incubated at 37°C overnight. After incubation, 4ml of NB was added to the plate, and the lysis area was evenly spread. The mixture was centrifuged at 15,000rpm for 5min, and the supernatant was filtered through a 0.22µm syringe filter and stored at 4°C.

### Phage titre determination

Phage concentrations were determined using a plaque assay adding 10µl of each 10-fold serial dilutions (from 10^-1^ to 10^-6^) of original phage stock. Plates were incubated at 37°C overnight, and plaques were counted to calculate plaque-forming units per millilitre (PFU/ml) using the formula:

Concentration = Number of plaques × 1000/10 × relevant dilution factor

### Bacteriophage lysis of target bacteria

To evaluate the lytic activity of individual phages and the phage cocktail, plate reader assays conducted in 12 well plates (greiner bio-one, Australia) were performed. Before conducting experiments, the optical density at 600nm (OD_600_) was measured for the content of each bacterial culture using a spectrophotometer (Biochrom Ltd., Cambridge, UK).

Experiments were conducted in two phases. Initially, single phage assays were conducted of three phages individually in the presence of different single bacterial strains, respectively. Two independent cultures of each bacterial strain were cultured, respectively in single phage assays. Phage cocktail assays were conducted in the presence of different single bacterial strains, respectively, or the different combinations of the three tested bacterial strains. Five independent cultures of each bacterial strain were cultured, respectively in phage cocktail to single bacterial growth assays. Two independent cultures of each bacterial strain were cultured and combined as different host combinations (Host combo A-E, listed in Table 2). After plate reader incubation in both phases, 1mL of the samples added with phage (Table 3) were collected to determine phage titre (Fig 1).

**Fig 1.**
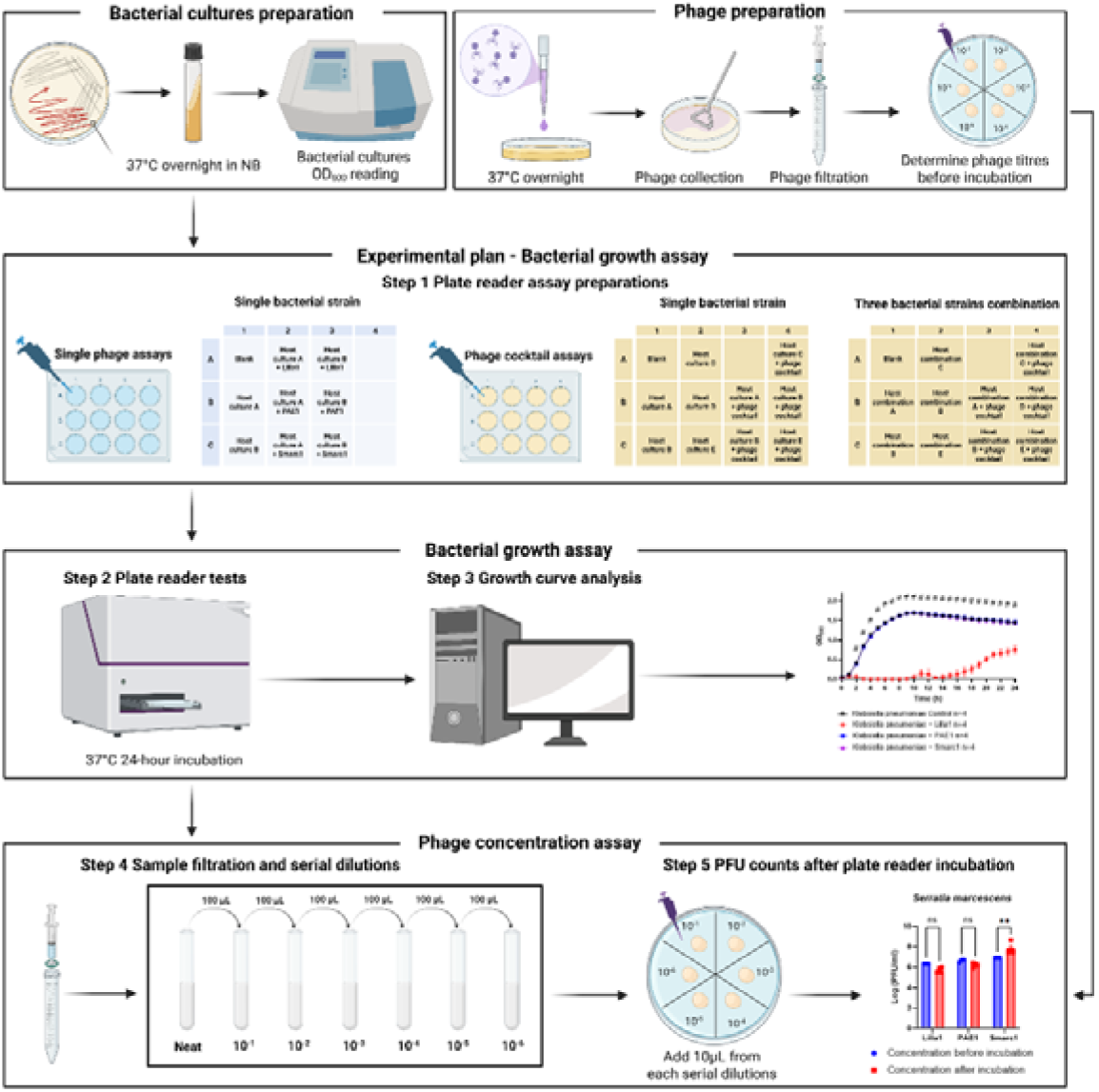
Experimental design to assess efficacy of Lilla1, PAE1, and Smarc1 on inhibiting host bacterial growth individually and in combination. The experimental design graph was illustrated by Biorender.com.

**Table 2.** The combination of host bacterial cultures used in phage cocktail assays.

| Name | Different combinations |
| --- | --- |
| Host combination A | *K. pneumoniae* culture A + *P. aeruginosa* culture A +<br>*S. marcescens* culture A |
| Host combination B | *K. pneumoniae* culture A + *P. aeruginosa* culture B +<br>*S. marcescens* culture A |
| Host combination C | *K. pneumoniae* culture A + *P. aeruginosa* culture A +<br>*S. marcescens* culture B |
| Host combination D | *K. pneumoniae* culture B + *P. aeruginosa* culture A +<br>*S. marcescens* culture A |
| Host combination E | *K. pneumoniae* culture B + *P. aeruginosa* culture B +<br>*S. marcescens* culture A |

**Table 3.** Samples collected for phage titre determination after plate reader assays.

| Assays | Samples collected |
| --- | --- |
| In single phage assay | Host culture A + Lilla1 |
|  | Host culture B + Lilla1 |
|  | Host culture A + PAE1 |
|  | Host culture B + PAE1 |
|  | Host culture A + Smarc1 |
|  | Host culture B + Smarc1 |
| In phage cocktail assay<br>(single bacterial strain tested) | Host culture A + phage cocktail |
|  | Host culture B + phage cocktail |
|  | Host culture C + phage cocktail |
|  | Host culture D + phage cocktail |
|  | Host culture E + phage cocktail |
| In phage cocktail assay<br>(host combination tested) | Host combo A + phage cocktail |
|  | Host combo B + phage cocktail |
|  | Host combo C + phage cocktail |
|  | Host combo D + phage cocktail |
|  | Host combo E + phage cocktail |

For each well, the final volume was set to 2000µL. To start with a consistent and low bacterial concentration, a starter OD_600_ reading of 0.05 was used. This starting value (equivalent to a low bacterial concentration) ensured that bacterial growth was clearly observed over time without reaching a stationary growth phase too early in the assay. The volume of overnight bacterial culture needed per well was calculated using the formula:

Volume of bacterial sample = (0.05 × 2000 µL) / OD_600_ of the overnight culture

The multiplicity of infection (MOI) was set at 1 for each phage. Each well was adjusted to 2000µL using NB. Blank wells contained only nutrient broth.

The OD_600_ of each well was automatically recorded every 60 minutes over a 24-hour period at 37°C with 200rpm meander corner well shaking using a microplate reader (CLARIOstar, BMGLabtech, Australia). Final OD_600_ values at each time point were calculated by subtracting the OD_600_ of the blank well from each raw reading:

Sample OD_600_ = Raw OD_600_ – Blank OD_600_

### Statistical analyses

Statistical analyses were conducted using GraphPad Prism software (version 10; GraphPad Software Inc., San Diego, California, USA). Group comparisons from the plate reader assay were performed using two-way ANOVA followed by Dunnett’s multiple comparisons test, and multiple comparisons were made at each timepoint. Phage concentration data were also analyzed using two-way ANOVA, followed by Tukey’s multiple-comparison test. Results are expressed as mean ± SEM, with statistical significance set at PlJ<lJ0.05.

## Results

### Effectiveness of Lilla1, PAE1, and Smarc1 in suppressing bacterial growth

To evaluate the specificity and efficacy of the individual phage Lilla1, PAE1, and Smarc1, we assessed their ability to inhibit the growth of their respective hosts (*Klebsiella pneumoniae*, *Pseudomonas aeruginosa*, and *Serratia marcescens*). Growth curves for each bacterial strain were recorded over 24h at 37°C using a CLARIOstar plate reader, following infection with both their respective cognate and other non-cognate phage. It was hypothesized that only phage specifically targeting designated bacteria would suppress bacterial growth of the target bacterium, whereas non-cognate phage would exhibit a growth pattern comparable to the non- infected bacteria.

Lilla1 demonstrated strong specificity in inhibiting growth of the target bacterium *K. pneumoniae* at 37°C. Lilla1 effectively suppressed *K. pneumoniae* growth during the first 10h of incubation. Despite some bacterial growth after this period, Lilla1 maintained significant inhibition of growth throughout the 24-h period. As expected, neither PAE1 nor Smarc1 impacted *K. pneumoniae* growth, similar to the non-phage infected control (Fig 2A). PAE1 phage exhibited high specificity in reducing via reducing peak growth levels. No statistical differences in *P. aeruginosa* growth, however, were observed between the control and phage-treated samples during the first 7h of incubation. The significant inhibition of *P. aeruginosa* growth by PAE1 was observed after an 8-hour incubation period. As expected, Lilla1 and Smarc1 did not impact *P. aeruginosa* growth (Fig 2B). Interestingly, a rapid and distinct inhibition pattern was observed for Smarc1 targeting of *S. marcescens* whereby the growth of *S. marcescens* was significantly reduced after incubation for only 2h at 37°C. Smarc1 delayed growth and reduced the peak growth level of *S. marcescens*. Although growth resumed after 13h, Smarc1-treated *S. marcescens* samples exhibited consistently lower maximum growth levels compared to untreated samples. Lilla1 and PAE1 had no effect on *S. marcescens* growth (Fig 2C).

**Fig 2.**
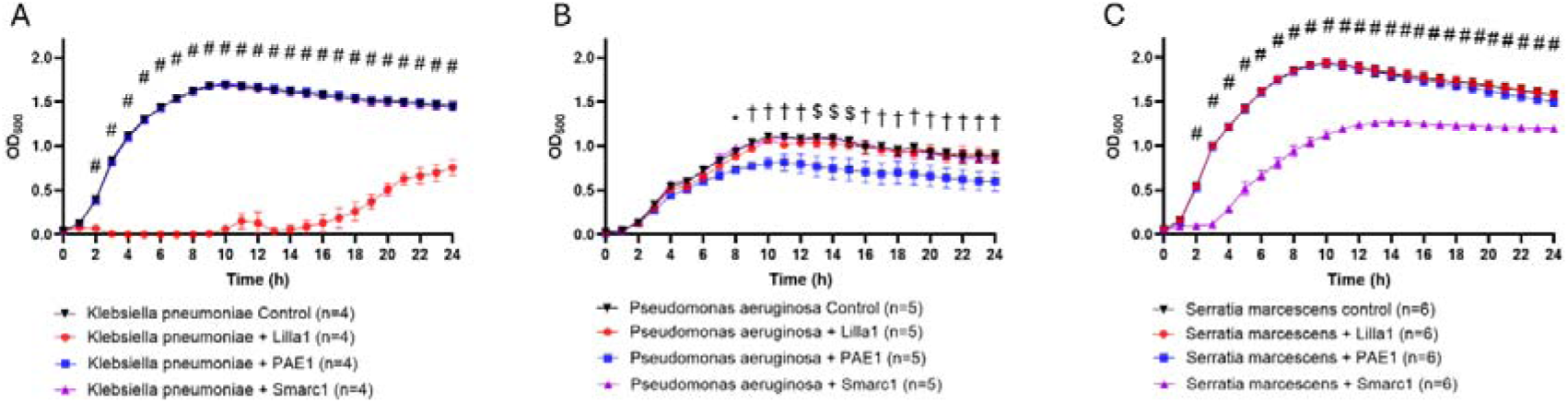
**Specific inhibition of *K. pneumoniae*, *P. aeruginosa* and *S. marcescens* by individual cognate phages; Lilla1, PAE1, and Smarc1.** Growth profiles for *K. pneumoniae*, *P. aeruginosa*, and *S. marcescens.* (A) *K. pneumoniae* infected with Lilla1, PAE1, or Smarc1; (B) *P. aeruginosa* infected with Lilla1, PAE1, or Smarc1; (C) *S. marcescens* infected with Lilla1, PAE1, or Smarc1. Two-way ANOVA was used to compare groups, with data presented as mean ± SEM, *p* < 0.05*\*, p < 0.01^†^, p < 0.001^$^, and p < 0.0001*^#^.

One notable observation was the regrowth of bacteria after an initial inhibition period, particularly when treated with their corresponding phage. This regrowth suggests the development of bacterial resistance to phage, potentially due to mechanisms such as mutations in phage receptor sites or the activation of bacterial defence systems.

The emergence of phage-resistant bacterial variants has been documented, highlighting the need for combination therapies and continued monitoring of bacterial response to phage infection [20].

### Host-specific inhibition patterns of individual phages Lilla1, PAE1, and Smarc1

Phage titration was measured before and after incubation with bacteria to determine the ability of phages to replicate while infecting host bacteria. An increase in phage concentration would indicate successful replication during bacterial targeting. In contrast, the concentration of non-specific phage is hypothesised to remain unchanged or decline. This profile and highlights the safety of phage therapy by demonstrating that phages lacking a specific host target do not replicate or persist in the environment, reducing the risk of unintended ecological impact.

As expected, the concentration of Lilla1 phage increased after a 24h incubation with its cognate host, *K. pneumoniae*. In contrast, when incubated with *K. pneumoniae*, the concentration of PAE1 phage remained stable post-incubation, indicating its inability to replicate in a non-cognate host. Smarc1 phage concentration decreased, however, suggesting that *K. pneumoniae* may have suppressed Smarc1 activity or contributed to its degradation (Fig 3A).

**Fig 3.**
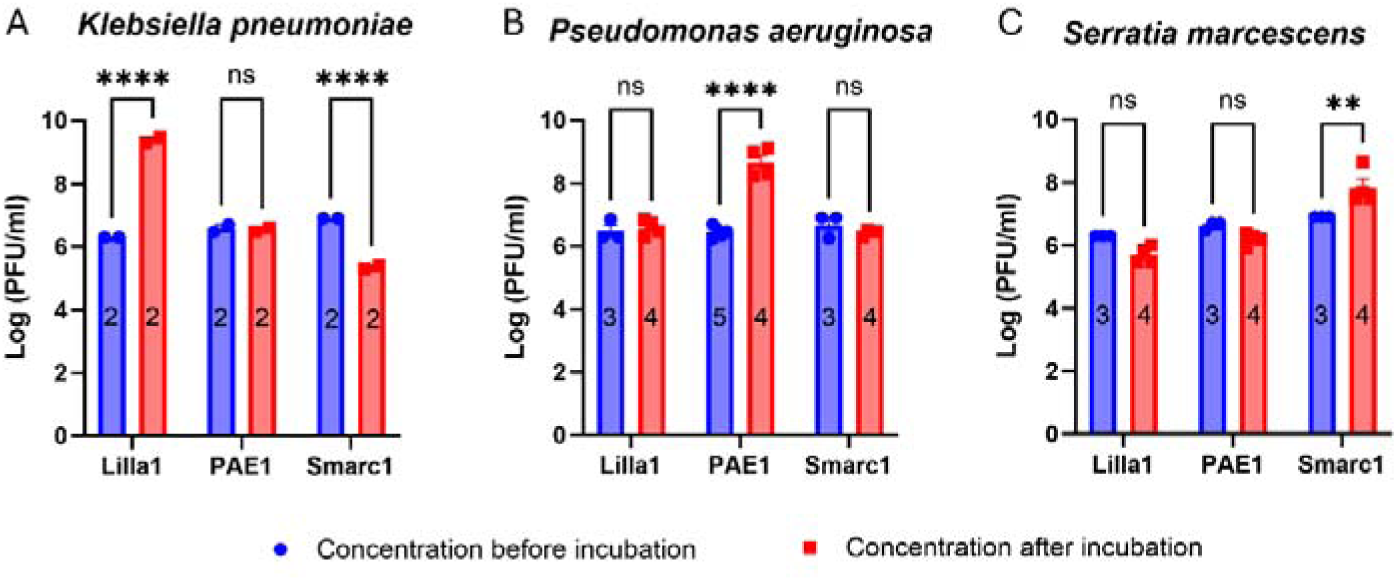
**Concentrations of individual phages before and after incubation with *K. pneumoniae*, *P. aeruginosa*, and *S. marcescens*, respectively.** Phage concentrations were measured before and after incubation to assess self-replication during bacterial infection. Phage concentration comparisons for Lilla1, PAE1, and Smarc1 before and after incubation with (A) *K. pneumoniae;* (B) *P. aeruginosa;* (C) *S. marcescens*. Two-way ANOVA was performed to analyze comparisons between groups. Data represented as mean ± SEM (*p* < 0.01 **, *p* < 0.0001 ****, *p* > 0.05, ns).

When incubated with *P. aeruginosa*, the cognate phage (i.e., PAE1) increased in concentration. Lilla1 concentration, however, remained unchanged during incubation with PAE1, confirming its inability to replicate in a non-cognate host. Interestingly, Smarc1 showed a slight but statistically insignificant decrease following incubation with PAE1, further supporting the notion that *P. aeruginosa* may suppress or degrade non-cognate phages (Fig 3B).

In *S. marcescens* bacterial cultures, the concentration of the cognate phage Smarc1 increased post-incubation, whereas Lilla1 and PAE1 showed no significant change (Fig 3C). Overall, these results highlight the specificity of Lilla1, PAE1, and Smarc1 for their respective hosts (i.e., *K. pneumoniae, P. aeruginosa,* and *S. marcescens*, respectively) as evidenced by bacterial growth inhibition and changes in phage concentration.

#### *In vitro* effectiveness of a bacteriophage cocktail against growth of K. pneumoniae, P. aeruginosa, and S. marcescens

A bacteriophage cocktail comprising Lilla1, PAE1, and Smarc1 at a 1:1:1 ratio was evaluated for efficacy in inhibiting growth of *K. pneumoniae*, *P. aeruginosa*, and *S. marcescens*, both individually and in combination. Bacterial growth was tracked using a 24-hour growth curve assay at 37°C with or without the phage cocktail.

Here we show that the phage cocktail (i.e., Lilla1, PAE1, and Smarc1) effectively suppressed growth of bacteria *K. pneumoniae*, *P. aeruginosa*, and *S. marcescens* both individually and as a combined bacterial group.

The phage cocktail (i.e., Lilla1, PAE1, and Smarc1) significantly inhibited *K. pneumoniae* growth such that growth was negligible during the first 8h of incubation. Although bacterial growth resumed after 8h, the cocktail continued to exert a strong inhibitory effect on growth (Fig 4A). When incubated with *P. aeruginosa*, the phage cocktail primarily reduced peak *P. aeruginosa* growth levels. No significant differences in *P. aeruginosa* growth were observed between control and phage-treated samples during the first 7 hours of incubation. However, after 8h, the phage cocktail significantly inhibited *P. aeruginosa* growth, demonstrating greater efficacy compared to PAE1 alone. (Fig 4B). The phage cocktail also showed a distinctive inhibition pattern against *S. marcescens*. Specifically, the cocktail delayed the onset of *S. marcescens* growth and significantly reduced peak growth. During the first 3h of the incubation period, cocktail-treated *S. marcescens* samples exhibited negligible growth compared to the control. The phage cocktail showed sustained inhibition of *S. marcescens* growth throughout the assay, with peak growth of phage-treated samples remaining significantly lower than controls (Fig 4C).

**Fig 4.**
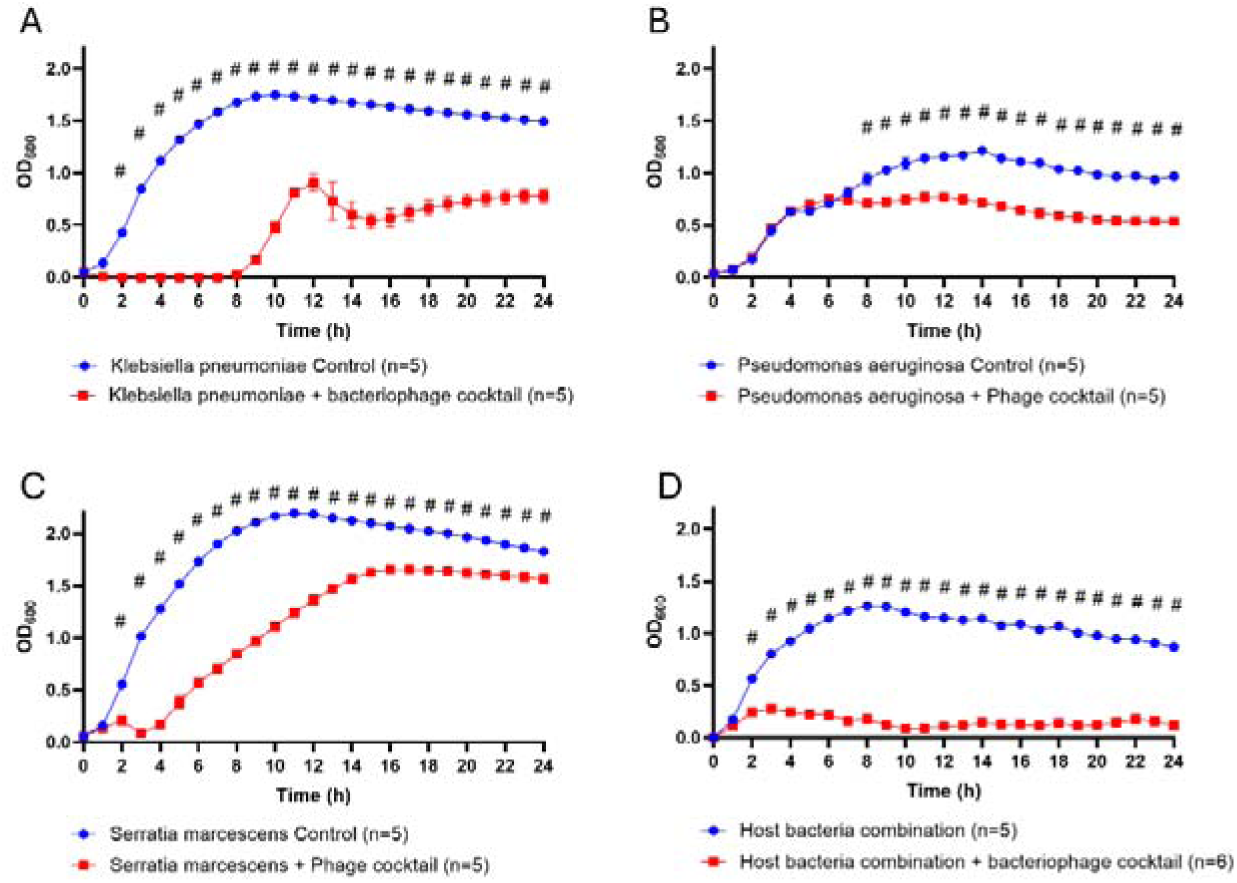
The Lilla1, PAE1 and Smarc1 bacteriophage cocktail inhibits growth of *K. pneumoniae*, *P. aeruginosa*, and *S. marcescens* individually and in combination. Growth profiles for *K. pneumoniae*, *PAO9503*, and *S. marcescens* and in combination with the Lilla1-PAE1-Smarc1 phage cocktail. (A) *K. pneumoniae* infected with Lilla1, PAE1, and Smarc1. (B) *P. aeruginosa* infected with Lilla1, PAE1, and Smarc1. (C) *S. marcescens* infected with Lilla1, PAE1, and Smarc1. (D) Combined pathogenic bacteria treated with the bacteriophage cocktail. Two-way ANOVA was used to analyze comparisons between groups. Data represented as mean ± SEM (p < 0.0001 (#); p > 0.05, ns).

Finally, when the efficacy of the cocktail against bacterial growth for the combination of *K. pneumoniae*, *P. aeruginosa*, and *S. marcescens* was evaluated the growth of mixed bacterial samples was continuously inhibited during the 24-h incubation period compared to control samples (Fig 4D).

### Evaluating phage cocktail self-replication and inhibition of *K. pneumoniae, P. aeruginosa,* and *S. marcescens*

Phage titrations were determined before and after exposure to host bacteria to assess the efficacy of the bacteriophage cocktail in inhibiting *K. pneumoniae*, *P. aeruginosa*, and *S. marcescens*, both individually and in combination. When phage cocktail was incubated with *K. pneumoniae*, only the cognate phage Lilla1 exhibited a significant increase in concentration, indicating its active replication when mixed with the host. The concentration of PAE1 remained unchanged after incubation with *K. pneumoniae*, whereas Smarc1 was decreased in concentration, further suggesting that *K. pneumoniae* may have suppressed Smarc1 activity or contributed to its degradation (Fig 5A). These results align with findings from our individual phage experiments, where Lilla1 consistently exhibited robust replication, PAE1 remained stable, whereas Smarc1 showed a less pronounced decrease in concentration. In the presence of *P. aeruginosa*, however, only the cognate phage (PAE1) exhibited an increase in concentration. In contrast, both Lilla1 and Smarc1 showed no change in concentration after incubation with *P. aeruginosa* (Fig 5B).

**Fig 5.**
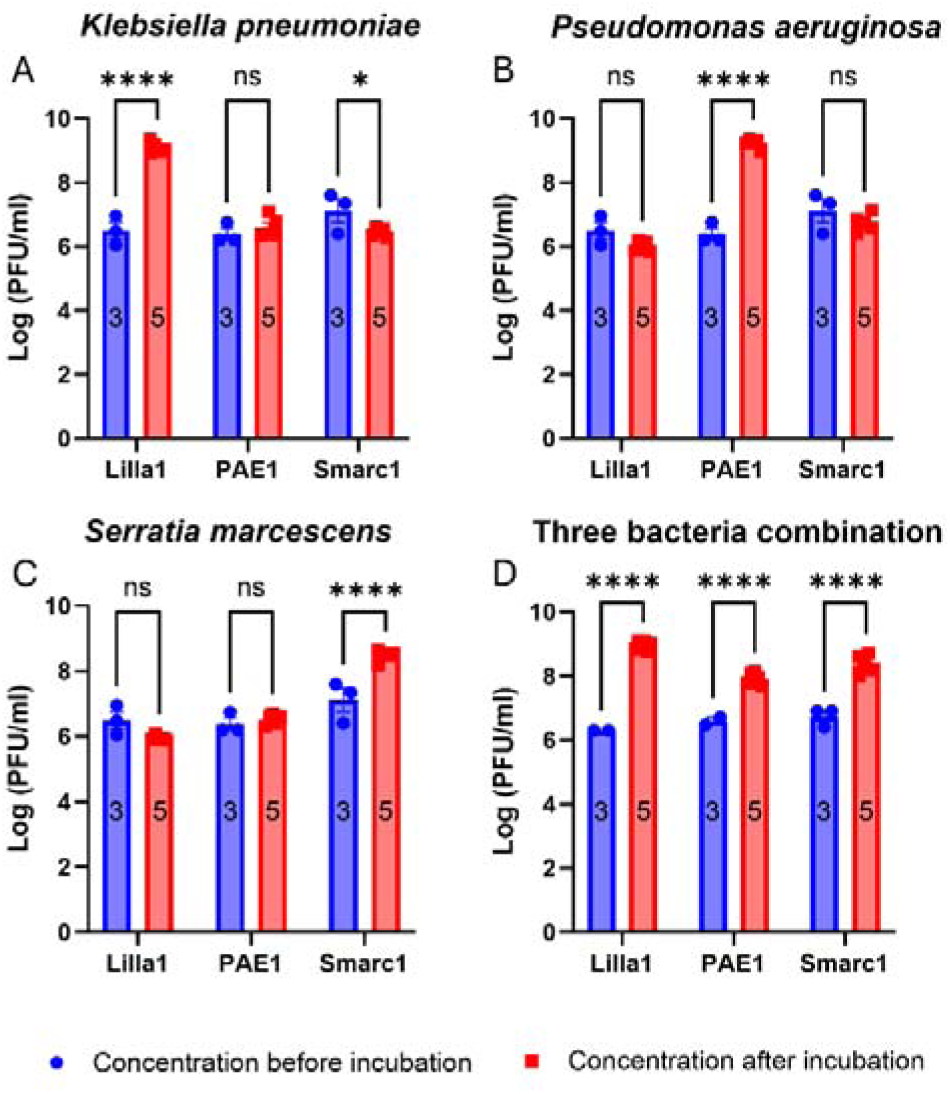
**Phage cocktail concentration before and after incubation with *K. pneumoniae*, *P. aeruginosa*, and *S. marcescens*, individually and in combination.** (A) Phage concentration comparisons between phage stock, before, and after incubation with *K. pneumoniae*. (B) Phage concentration comparisons between phage stock, before, and after incubation with *P. aeruginosa*. (C) Phage concentration comparisons between phage stock, before, and after incubation with *S. marcescens*. (D) Phage concentration comparisons between phage stock, before, and after incubation with the combination of *K. pneumoniae*, *P. aeruginosa*, and *S. marcescens*. Two-way ANOVA was used to compare the groups. Data are represented as mean ± SEM; p<0.05 *; p<0.0001****; p>0.05, ns.

When targeting *S. marcescens*, only the cognate phage (Smarc1) increased in concentration. The concentration of PAE1 remained unchanged after incubation with *S. marcescens*, whereas the concentration of Lilla1 decreased, suggesting that *S. marcescens* may have suppressed Lilla1 and PAE1 activity or contributed to the degradation of Lilla1 (Fig 5C). These results align with individual phage assays against *S. marcescens*. Interestingly, Smarc1 displayed enhanced efficacy in the cocktail, as evidenced by a more robust inhibition of *S. marcescens* growth with higher statistical significance compared to its performance in single-phage assays. When the phage cocktail was applied to target *K. pneumoniae*, *P. aeruginosa*, and *S. marcescens* in combination, the concentrations of all three phage significantly increased compared to pre-incubation levels (Fig 5D), demonstrating effective targeting of all three hosts. These results suggest that combining the phages in a cocktail not only maintains individual efficacy but also enhances the replication and activity of Smarc1, potentially offering a synergistic advantage for controlling multi-bacterial infections.

The phage cocktail demonstrated robust inhibition of *K. pneumoniae*, *P. aeruginosa*, and *S. marcescens* in combination. In the presence of the phage cocktail, bacterial growth remained minimal throughout a 24-hour incubation period, and each phage within the cocktail showed a significant increase in concentration. These findings indicate effective targeting and amplification in response to the respective bacterial hosts. These results highlight potential advantages of using a phage cocktail to combat complex bacterial infections, especially in cases where multiple pathogens are implicated in GI dysfunction.

## Discussion

We investigated the effects of phage cocktail treatment including Lilla1, PAE1, and Smarc1 on the targeted bacterial growth of *K. pneumoniae*, *P. aeruginosa*, and *S. marcescens in vitro*, and potential implications for therapeutic applications. Phage treatment significantly inhibited the growth of bacterial populations over time, consistent with previous studies exploring phage therapy as an alternative to antibiotics [24, 25]. Bacterial reduction occurred in a host-dependent manner, indicating target specificity of these phages. However, variations in bacterial regrowth observed in some cases suggest potential bacterial resistance potentially due to mechanisms involving CRISPR-Cas systems or mutations in phage receptor sites [26, 27]. Future studies should investigate the genetic basis of these resistance patterns and explore strategies to overcome them, such as phage cocktail formulations or the use of phage-derived enzymes [28–30].

A key finding in our study was the observation of a biphasic response pattern during phage therapy, characterized by an initial rapid decline in bacterial populations followed by partial regrowth over time. This phenomenon suggests a dynamic interaction between bacterial defence mechanisms and phage infectivity. The biphasic response pattern we observed is consistent with previous studies reporting initial bacterial decline followed by regrowth due to resistance or persistent cell survival [31, 32]. This phenomenon presents a significant challenge for phage therapy as it limits long-term bacterial suppression and may contribute to the development of phage resistance. To address this challenge, future research should explore combination therapies involving phage and antibiotics to prevent resistance emergence and enhance bacterial eradication [25, 33].

Our findings provide valuable insights into phage-bacteria interactions; however, certain limitations should be acknowledged. *In vitro* models, though useful for controlled investigations, do not fully replicate the complexity of *in vivo* environments, where factors such as immune responses and host-pathogen interactions significantly influence phage efficacy [34]. Future research should incorporate *in vivo* models to validate the therapeutic potential of phage in physiologically relevant conditions.

## Conclusions

In summary, our findings show that *K. pneumoniae* phage Lilla1, *P. aeruginosa* phage PAE1, and *S. marcescens* phage Smarc1 effectively inhibit their respective host bacteria and self-replicate during infection. The phage cocktail exhibited a synergistic effect, providing continuous suppression of bacterial growth when applied to a mixture of pathogenic bacteria. Given that previous studies report that gram-negative bacteria, such as those studied here, can increase gut permeability, our findings that Lilla1, PAE1, and Smarc1 effectively suppress bacterial growth and limit bacterial proliferation suggest that further development of this phage cocktail could offer a therapeutic approach for treating GI dysfunctions associated with pathogenic bacterial overgrowth. However, bacterial resistance remains a key challenge, underscoring the need for continued research on resistance mechanisms and therapeutic optimization. By addressing these challenges, phage therapy may emerge as a viable alternative to conventional antibiotics in the fight against bacterial infections.

## Acknowledgements

We thank Associate Professor Steve Petrovski at La Trobe University for generously donating all bacterial and phage strains. We also thank Professor Ashley Franks, Associate Professor Steve Petrovski, and Professor Elisa Hill-Yardin for reviewing and editing the original manuscript.

## Author Contributions

Yuansong Li (Investigation [Lead], Data analysis [Lead], Data curation [equal], Writing – original draft [equal]), Ashley Franks (Conceptualization [equal], Project administration [equal], Supervision [equal]), Elisa Hill-Yardin (Conceptualization [equal], Resources [Lead], Project administration [equal], Validation [equal], Supervision [equal], Writing – review & editing [equal]), Steve Petrovski (Conceptualization [Lead], Resources [Lead], Project administration [equal], Validation [equal], Data curation [equal], Supervision [equal], Writing – review & editing [equal],)

## Funding

This study was supported by National Health and Medical Research Council (NHMRC) (Application No. APP2003848).

## Conflict of interest

The authors declare no conflict of interest.

## Data availability

The data underlying this article will be shared on reasonable request to the corresponding author.

## Notes

### Competing Interest Statement

The authors have declared no competing interest.

## References

1. Ochieng JB, Boisen N, Lindsay B, Santiago A, Ouma C, Ombok M, et al. Serratia marcescens is injurious to intestinal epithelial cells. Gut Microbes. 2014;5(6):729–36. Epub 2014/11/27. doi: 10.4161/19490976.2014.972223. PubMed PMID: 25426769; PubMed Central PMCID: PMCPMC4615285.

2. Stephens M, von der Weid PY. Lipopolysaccharides modulate intestinal epithelial permeability and inflammation in a species-specific manner. Gut Microbes. 2020;11(3):421–32. Epub 2019/06/18. doi: 10.1080/19490976.2019.1629235. PubMed PMID: 31203717; PubMed Central PMCID: PMCPMC7524286.

3. Bischoff SC, Barbara G, Buurman W, Ockhuizen T, Schulzke JD, Serino M, et al. Intestinal permeability--a new target for disease prevention and therapy. BMC Gastroenterol. 2014;14:189. Epub 2014/11/20. doi: 10.1186/s12876-014-0189-7. PubMed PMID: 25407511; PubMed Central PMCID: PMCPMC4253991.

4. Farre R, Fiorani M, Rahiman SA, Matteoli G. Intestinal Permeability, Inflammation and the Role of Nutrients. Nutrients. 2020;12(4). doi: ARTN 1185 10.3390/nu12041185. PubMed PMID: WOS:000531831300305.

5. Brandtzaeg P. The gut as communicator between environment and host: Immunological consequences. Eur J Pharmacol. 2011;668:S16-S32. doi: 10.1016/j.ejphar.2011.07.006. PubMed PMID: WOS:000296599700004.

6. Podschun R, Ullmann U. Klebsiella spp. as nosocomial pathogens: epidemiology, taxonomy, typing methods, and pathogenicity factors. Clin Microbiol Rev. 1998;11(4):589–603. Epub 1998/10/10. doi: 10.1128/CMR.11.4.589. PubMed PMID: 9767057; PubMed Central PMCID: PMCPMC88898.

7. Gellatly SL, Hancock RE. Pseudomonas aeruginosa: new insights into pathogenesis and host defenses. Pathog Dis. 2013;67(3):159–73. Epub 2013/04/27. doi: 10.1111/2049-632X.12033. PubMed PMID: 23620179.

8. Hejazi A, Falkiner FR. Serratia marcescens. J Med Microbiol. 1997;46(11):903-12. Epub 1997/11/22. doi: 10.1099/00222615-46-11-903. PubMed PMID: 9368530.

9. Russo TA, Marr CM. Hypervirulent Klebsiella pneumoniae. Clin Microbiol Rev. 2019;32(3). Epub 2019/05/17. doi: 10.1128/CMR.00001-19. PubMed PMID: 31092506; PubMed Central PMCID: PMCPMC6589860.

10. Spiers AJ, Buckling A, Rainey PB. The causes of Pseudomonas diversity. Microbiology (Reading). 2000;146 (Pt 10):2345–50. Epub 2000/10/06. doi: 10.1099/00221287-146-10-2345. PubMed PMID: 11021911.

11. Mahlen SD. Serratia infections: from military experiments to current practice. Clin Microbiol Rev. 2011;24(4):755–91. Epub 2011/10/07. doi: 10.1128/CMR.00017-11. PubMed PMID: 21976608; PubMed Central PMCID: PMCPMC3194826.

12. Sandler RH, Finegold SM, Bolte ER, Buchanan CP, Maxwell AP, Vaisanen ML, et al. Short-term benefit from oral vancomycin treatment of regressive-onset autism. J Child Neurol. 2000;15(7):429–35. Epub 2000/08/02. doi: 10.1177/088307380001500701. PubMed PMID: 10921511.

13. Grossi E, Melli S, Dunca D, Terruzzi V. Unexpected improvement in core autism spectrum disorder symptoms after long-term treatment with probiotics. SAGE Open Med Case Rep. 2016;4:2050313X16666231. Epub 2016/09/14. doi: 10.1177/2050313X16666231. PubMed PMID: 27621806; PubMed Central PMCID: PMCPMC5006292.

14. Kaluzna-Czaplinska J, Blaszczyk S. The level of arabinitol in autistic children after probiotic therapy. Nutrition. 2012;28(2):124–6. Epub 2011/11/15. doi: 10.1016/j.nut.2011.08.002. PubMed PMID: 22079796.

15. Kang DW, Adams JB, Gregory AC, Borody T, Chittick L, Fasano A, et al. Microbiota Transfer Therapy alters gut ecosystem and improves gastrointestinal and autism symptoms: an open-label study. Microbiome. 2017;5(1):10. Epub 2017/01/27. doi: 10.1186/s40168-016-0225-7. PubMed PMID: 28122648; PubMed Central PMCID: PMCPMC5264285.

16. Ventola CL. The antibiotic resistance crisis: part 2: management strategies and new agents. P T. 2015;40(5):344–52. Epub 2015/05/20. PubMed PMID: 25987823; PubMed Central PMCID: PMCPMC4422635.

17. Gareau MG, Sherman PM, Walker WA. Probiotics and the gut microbiota in intestinal health and disease. Nat Rev Gastroenterol Hepatol. 2010;7(9):503–14. Epub 2010/07/29. doi: 10.1038/nrgastro.2010.117. PubMed PMID: 20664519; PubMed Central PMCID: PMCPMC4748966.

18. Wang JG, Liang Q, Dou HH, Ou Y. The global incidence of adverse events associated with fecal microbiota transplantation in children over the past 20 years: A systematic review and meta-analysis. J Gastroenterol Hepatol. 2022;37(11):2031–8. Epub 2022/09/07. doi: 10.1111/jgh.15996. PubMed PMID: 36066910.

19. Wang S, Xu M, Wang W, Cao X, Piao M, Khan S, et al. Systematic Review: Adverse Events of Fecal Microbiota Transplantation. PLoS One. 2016;11(8):e0161174. Epub 2016/08/17. doi: 10.1371/journal.pone.0161174. PubMed PMID: 27529553; PubMed Central PMCID: PMCPMC4986962.

20. Pourraz CBJ. Phage therapy as a potential solution in the fight against AMR: obstacles and possible futures. Palgrave Communications. 2020;6. doi: 10.1057/s41599-020-0478-4.

21. Sulakvelidze A, Alavidze Z, Morris JG, Jr. Bacteriophage therapy. Antimicrob Agents Chemother. 2001;45(3):649–59. Epub 2001/02/22. doi: 10.1128/AAC.45.3.649-659.2001. PubMed PMID: 11181338; PubMed Central PMCID: PMCPMC90351.

22. Barr JJ. A bacteriophages journey through the human body. Immunol Rev. 2017;279(1):106–22. Epub 2017/09/01. doi: 10.1111/imr.12565. PubMed PMID:

23. 28856733.

23. Barr JJ, Auro R, Sam-Soon N, Kassegne S, Peters G, Bonilla N, et al. Subdiffusive motion of bacteriophage in mucosal surfaces increases the frequency of bacterial encounters. Proc Natl Acad Sci U S A. 2015;112(44):13675–80. Epub 2015/10/21. doi: 10.1073/pnas.1508355112. PubMed PMID: 26483471; PubMed Central PMCID: PMCPMC4640763.

24. Niu YD, Liu H, Du H, Meng R, Sayed Mahmoud E, Wang G, et al. Efficacy of Individual Bacteriophages Does Not Predict Efficacy of Bacteriophage Cocktails for Control of Escherichia coli O157. Front Microbiol. 2021;12:616712. Epub 2021/03/16. doi: 10.3389/fmicb.2021.616712. PubMed PMID: 33717006; PubMed Central PMCID: PMCPMC7943454.

25. Cui L, Watanabe S, Miyanaga K, Kiga K, Sasahara T, Aiba Y, et al. A Comprehensive Review on Phage Therapy and Phage-Based Drug Development. Antibiotics (Basel). 2024;13(9). Epub 2024/09/28 22:43. doi: 10.3390/antibiotics13090870. PubMed PMID: 39335043; PubMed Central PMCID: PMCPMC11428490.

26. Dimitriu T, Kurilovich E, Lapinska U, Severinov K, Pagliara S, Szczelkun MD, et al. Bacteriostatic antibiotics promote CRISPR-Cas adaptive immunity by enabling increased spacer acquisition. Cell Host Microbe. 2022;30(1):31–40 e5. Epub 2021/12/22. doi: 10.1016/j.chom.2021.11.014. PubMed PMID: 34932986.

27. Li N, Zeng Y, Wang M, Bao R, Chen Y, Li X, et al. Characterization of Phage Resistance and Their Impacts on Bacterial Fitness in Pseudomonas aeruginosa. Microbiol Spectr. 2022;10(5):e0207222. Epub 2022/09/22. doi: 10.1128/spectrum.02072-22. PubMed PMID: 36129287; PubMed Central PMCID: PMCPMC9603268.

28. Hussain FA, Dubert J, Elsherbini J, Murphy M, VanInsberghe D, Arevalo P, et al. Rapid evolutionary turnover of mobile genetic elements drives bacterial resistance to phages. Science. 2021;374(6566):488-92. Epub 2021/10/22. doi: 10.1126/science.abb1083. PubMed PMID: 34672730.

29. Subramanian A. Emerging roles of bacteriophage-based therapeutics in combating antibiotic resistance. Front Microbiol. 2024;15:1384164. Epub 2024/07/22. doi: 10.3389/fmicb.2024.1384164. PubMed PMID: 39035437; PubMed Central PMCID: PMCPMC11257900.

30. Gordillo Altamirano FL, Barr JJ. Phage Therapy in the Postantibiotic Era. Clin Microbiol Rev. 2019;32(2). Epub 2019/01/18. doi: 10.1128/CMR.00066-18. PubMed PMID: 30651225; PubMed Central PMCID: PMCPMC6431132.

31. Kunnath AP, Suodha Suoodh M, Chellappan DK, Chellian J, Palaniveloo K. Bacterial Persister Cells and Development of Antibiotic Resistance in Chronic Infections: An Update. Br J Biomed Sci. 2024;81:12958. Epub 2024/08/22. doi: 10.3389/bjbs.2024.12958. PubMed PMID: 39170669; PubMed Central PMCID: PMCPMC11335562.

32. Wood TK, Knabel SJ, Kwan BW. Bacterial persister cell formation and dormancy. Appl Environ Microbiol. 2013;79(23):7116–21. Epub 2013/09/17. doi: 10.1128/AEM.02636-13. PubMed PMID: 24038684; PubMed Central PMCID: PMCPMC3837759.

33. Gordillo Altamirano FL, Kostoulias X, Subedi D, Korneev D, Peleg AY, Barr JJ. Phage-antibiotic combination is a superior treatment against Acinetobacter baumannii in a preclinical study. EBioMedicine. 2022;80:104045. Epub 2022/05/11. doi: 10.1016/j.ebiom.2022.104045. PubMed PMID: 35537278; PubMed Central PMCID: PMCPMC9097682.

34. Hernandez CA, Koskella B. Phage resistance evolution in vitro is not reflective of in vivo outcome in a plant-bacteria-phage system. Evolution. 2019;73(12):2461–75. Epub 2019/08/23. doi: 10.1111/evo.13833. PubMed PMID: 31433508.

